# The spatial self-organization within pluripotent stem cell colonies is continued in detaching aggregates

**DOI:** 10.1101/2020.11.03.366518

**Authors:** Mohamed H. Elsafi Mabrouk, Roman Goetzke, Giulio Abagnale, Burcu Yesilyurt, Lucia Salz, Kira Zeevaert, Zhiyao Ma, Marcelo A. S. Toledo, Ronghui Li, Ivan G. Costa, Vivek Pachauri, Uwe Schnakenberg, Martin Zenke, Wolfgang Wagner

## Abstract

Colonies of induced pluripotent stem cells (iPSCs) reveal aspects of self-organization even under culture conditions that maintain pluripotency. To investigate the dynamics of this process under spatial confinement, we used either polydimethylsiloxane (PDMS) pillars or micro-contact printing of vitronectin. There was a progressive upregulation of OCT4, E-cadherin, and NANOG within 70 µm from the outer rim of iPSC colonies. Single-cell RNA-sequencing demonstrated that *OCT4*^high^ subsets have pronounced up-regulation of the TGF-β pathway, particularly of NODAL and its inhibitor LEFTY, at the rim of the colonies. Furthermore, calcium-dependent cell-cell interactions were found to be relevant for the self-organization. Interestingly, after 5 to 7 days, the iPSC colonies detached spontaneously from micro-contact printed substrates to form 3D aggregates. This new method allowed generation of embryoid bodies (EBs) of controlled size, without any enzymatic or mechanical treatment. Within the early 3D aggregates, the radial organization and differential gene expression continued in analogy to the changes observed during self-organization of iPSC colonies. Our results provide further insight into the gradual self-organization within iPSC colonies and at their transition into EBs.

## Introduction

Colonies of pluripotent stem cells (PSCs) are not homogeneous, but comprise subpopulations that express different levels of pluripotency markers (Nguyen et al., 2018; Singh et al., 2007; Stewart et al., 2006). These states are interconvertible: when isolated and re-plated, all subpopulations re-express the whole spectrum of pluripotency markers found in the original culture (Hough et al., 2009; Lau et al., 2020). Higher expression of pluripotency markers is often observed in the subpopulation localized at the border of colonies, which coincides with higher capacity of colony formation and self-renewal ability (Hough et al., 2014; Warmflash et al., 2014). This was consistently observed with various approaches for spatial confinement to control the size and shape of PSC colonies (Deglincerti et al., 2016).

The self-organization within PSC colonies may recapitulate some aspects of early embryogenesis, but the underlying mechanisms and dynamics are not sufficiently understood. Various models have been proposed for this self-organization, including differential activation of the transforming growth factor beta (TGF-ß) pathway between colony edge and center (Warmflash et al., 2014), which might be modulated by Yes-associated protein (YAP) and Transcriptional coactivator with PDS binding motif (TAZ) activity (Narimatsu et al., 2015; Varelas et al., 2010), ii) inaccessibility of receptors at the center of the colony (Etoc et al., 2016; Nallet-Staub et al., 2015), or iii) a Turing pattern with reaction-diffusion of activators and inhibitors (Tewary et al., 2017). Furthermore, it remains unclear how the spatial heterogeneity within colonies is reflected in transcriptomes of individual cells, or upon transition to three-dimensional organization (3D) towards embryoid bodies (EBs). In this study, we investigated the dynamics of self-organization of induced pluripotent stem cells (iPSCs) in spatially confined two-dimensional (2D) colonies and in self-detached 3D aggregates.

## Results

### Progressive self-organization of pluripotency factors in confined colonies

Colonies of iPSCs revealed marked up-regulation of OCT4, E-cadherin, and NANOG at their outer rim region when cultured for six days, e.g. on flat polydimethylsiloxane (PDMS; Fig. 1A). To systematically investigate the impact of culture time and colony size on this self-organization, we utilized circular PDMS micro-pillars of 200 µm height with diameters of 400, 600, and 800 µm (Suppl. Fig. S1A). This setup enables spatially confined cell growth on top of the pillars. Over six days, we observed marked and progressive up-regulation of OCT4, E-cadherin, and NANOG within a ring-shaped region of about 50 to 70 µm from the colony border, irrespective of pillar size (Fig.1B). The same increasing up-regulation from the rim was also observed for colonies on elliptical pillars with equivalent growth areas (Suppl. Fig. S1B). Imaging and handling of PDMS micro-pillars was cumbersome and molecular analysis hampered by cell growth between the pillars. Therefore, we alternatively used the array of pillars as a stamp for micro-contact printing (µCP) of the adhesion protein vitronectin on either tissue culture plastic (TCP) or glass substrates (Suppl. Fig. S1C). When iPSCs were seeded on these µCP substrates, their growth was also restricted to circular areas (Suppl. Fig. S1D) and we observed a very similar up-regulation of OCT4 and E-cadherin at the outer region, which was progressive over six days - again within about 50 to 70 µm distance from the border of the colony (Fig.1C-D; Suppl. Fig. S1E).

**Figure 1:**
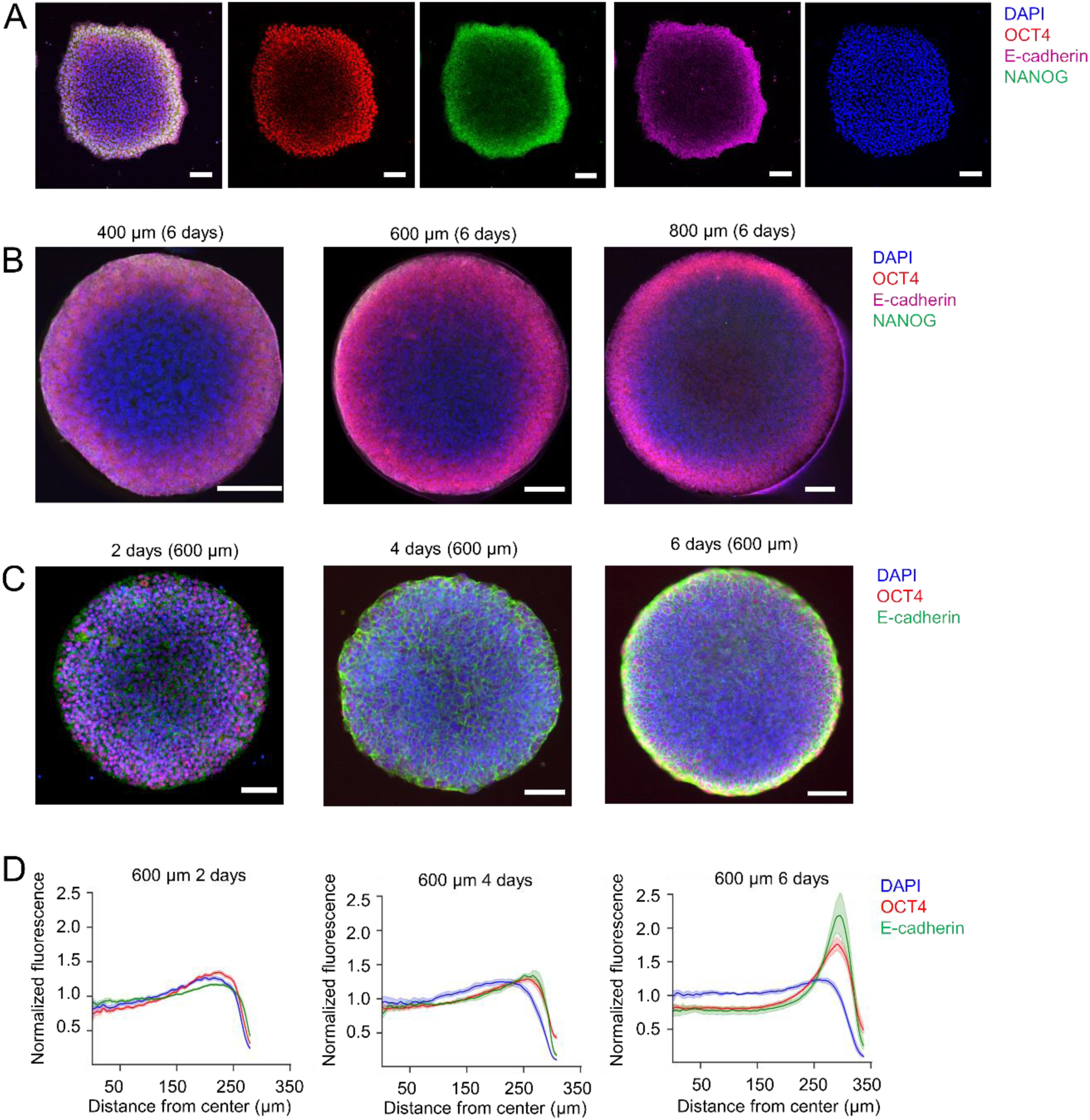
Spatial self-organization within iPSCs colonies. **A)** Higher expression of pluripotency markers (OCT4, E-cadherin, and NANOG) is observed at the border regions of induced pluripotent stem cell (iPSC) colonies, which were cultured on flat PDMS substrates for 6 days (scale bar: 50 µm). **B)** Up-regulation of NANOG, OCT4, and E-cadherin was particularly observed at the outer region of spatially confined iPSC colonies on PDMS micro-pillars with different diameters (Ø 400 µm, 600 µm, or 800 µm) after 6 days (scale bar: 100 µm). **C)** Continuous self-organization was observed on spatially confined iPSC colonies upon micro-contact printing (µCP) of vitronectin on tissue culture plastic (at d2, d4, and d6; scale bar: 100 µm). **D)** Quantification of immunofluorescence signals of iPSC colonies on µCP substrates demonstrate progressive organization (in analogy to C; n = 15 per time point).

### Gene expression is heterogeneous in spatially organized colonies

We anticipated that the spatial self-organization within iPSC colonies was also reflected on transcriptomic level and that this might shed light into the underlying mechanism. Therefore, we performed single-cell RNA sequencing (scRNA-seq) after 5 days of culture on 600 µm diameter µCP substrates using the 10x genomics platform. As a surrogate for the spatial organization of individual cells, we stratified the cells by their *POU5F1* (OCT4) expression level: in t-distributed stochastic neighbor embedding (t-SNE) plots the 1,000 cells with highest and lowest *OCT4* expression clustered with a smooth transition (Fig. 2A). When we directly compared these two subpopulations, there were 151 differentially expressed genes (fold change > ln1.5 and adjusted p-value < 0.05; Suppl. Table S1). Interestingly, *NODAL, LEFTY1*, and *LEFTY2* were the highest up-regulated genes in the *OCT4*^high^ subset (fold-change = 6.2, 5.45, and 4.74, respectively; Fig. 2B). These cytokines of the TGF-β superfamily have important functions in pluripotency maintenance as well as cellular differentiation in early embryogenesis. *OCT4*^high^ cells showed also up-regulation of other pluripotency markers, such as *NANOG, ZSCAN2, and MYC*, and of genes belonging to pluripotency maintenance pathways, such as *DUSP6* (FGF2/ERK pathway) and *SKIL* (TGF-ß pathway) (Fig. 2C). Very similar results were observed with an independent biological replicate of another iPSC-line (Suppl. Fig. S2A,B). Gene set enrichment analysis (GSEA) revealed that particularly genes of the TGF-ß signaling pathway are enriched in the *OCT4*^high^ subset (Suppl. Fig. S2C). Notably, there was a striking similarity in differential gene expression between the *OCT4*^high^ and *OCT4*^low^ subsets with a recently published study that compared subsets of embryonic stem cells (ESCs) with higher and lower self-renewal capacity (GCTM2^high^ CD9^high^ EPCAM^high^ *versus* GCTM2^low^ CD9^low^ EPCAM^low^) (Lau et al., 2020) (Suppl. Fig. S2D). Overall, the results suggested that the *OCT4*^high^ cells in the border region are in a more primitive state and that TGF-β pathways might be involved in its regulation.

**Figure 2:**
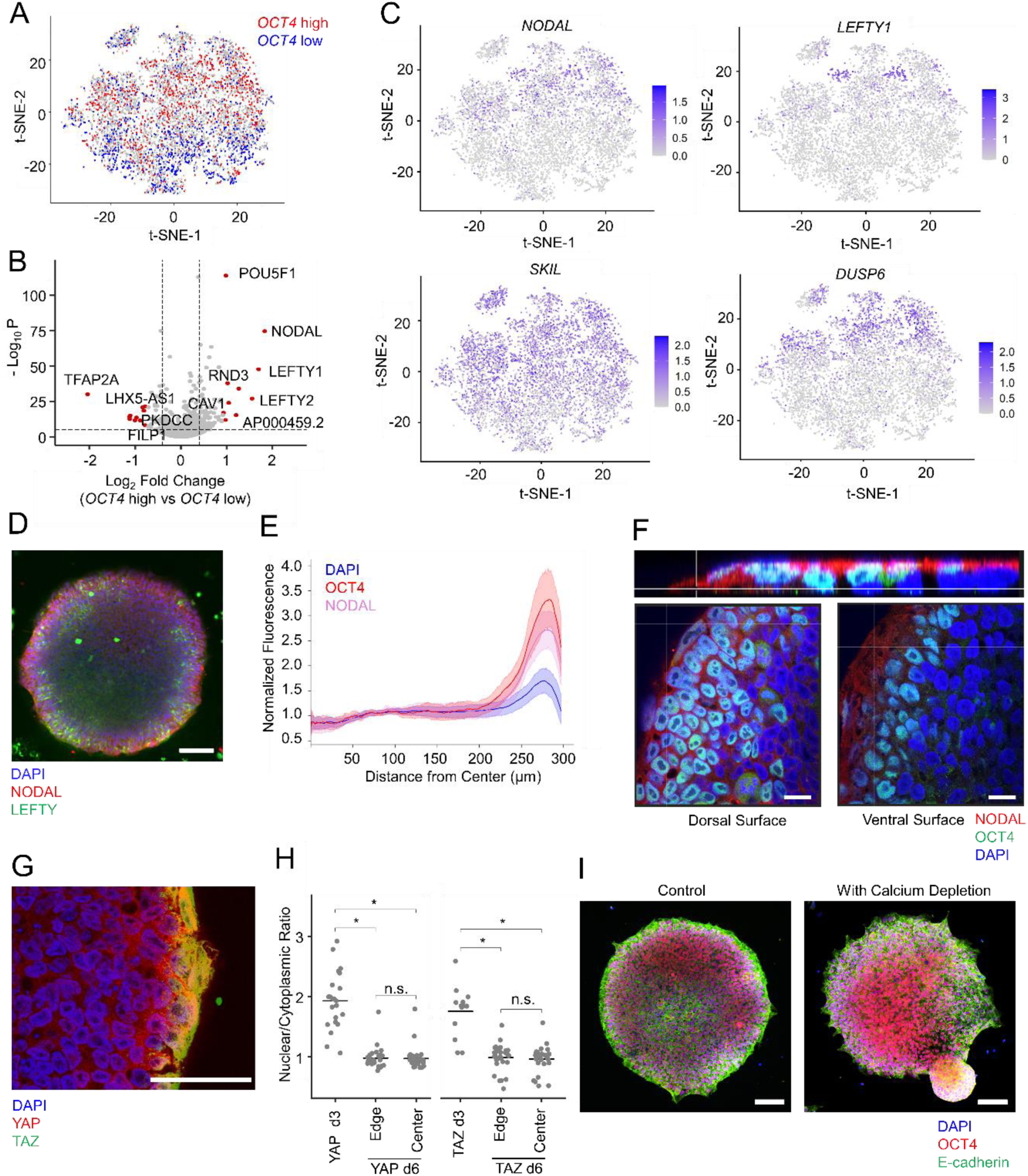
Relevant parameters for the spatial self-organization of iPSC colonies. **A)** Single-cell RNA-sequencing was performed with self-organized iPSC colonies at day 5 on µCP substrates (Ø 600 µm). The t-SNE plot highlights *OCT4*^high^ (red) and *OCT4*^low^ (blue) subfractions. **B)** Volcano plot of differential gene expression in *OCT4*^high^ *versus OCT4*^Low^ subsets demonstrates that *NODAL, LEFTY1*, and *LEFTY2* have highest up-regulation in *OCT4*^high^ subsets. **C)** t-SNE plots depict the heterogeneity of normalized expression of *NODAL, LEFTY1, SKIL, DUSP6* within the same cell preparations. **D)** Immunofluorescence image of iPSC at day 6 on µCP substrate (Ø 600 µm) demonstrates that expression of NODAL has similar spatial organization as OCT4, whereas LEFTY reveals rather patchy expression at the border (scale bar: 100 µm). **E)** Quantification of the radial profile of immunofluorescence signals of OCT4 and NODAL (n = 20 colonies). **F)** Perpendicular cut of Z-stack of spatially confined colony showing that on the dorsal surface of the colonies there is more expression of NODAL compared to the ventral surface of a self-organized colony (day 6 on µCP substrate; Ø 600 µm; scale bar: 20 µm). **G)** YAP and TAZ staining of a spatially confined iPSC colony (day 6 on µCP substrate; Ø 600 µm) demonstrates up-regulation of both proteins at the edge of the colony (scale bar: 50 µm). **H)** Quantification of nuclear/cytoplasmic ration of YAP and TAZ within spatially confined colonies at day 3 or day 6 (either center or edge of the colony; on µCP substrate, Ø 600 µm) demonstrates shift to the nuclear compartment at later time points (12 colonies for d3, 25 colonies for d6; \**P* < .001; n.s. = not significant; t-test with Bonferroni correction). **I)** Immunofluorescence image depicts spatial organization of OCT4 and E-cadherin (on µCP substrates, Ø 600 µm, d6) without or with intermittent calcium depletion by 2 mM EGTA (for 20 minutes at d5; scale bar: 50 µm).

In fact, confocal immunofluorescence analysis of NODAL and LEFTY showed similar up-regulation at the border region of confined iPSC colonies with continuous progression over six days, similar to the segregation of OCT4 (Fig. 2D, 2E). While NODAL was homogeneously up-regulated in the *OCT4*^high^ cells, LEFTY staining revealed a patchy expression within *OCT4*^high^ cells. This is in line with the clustered LEFTY expression in the t-SNE plots of scRNA-Seq data (Fig. 2C). Furthermore, NODAL staining was more prominent on the dorsal surface of the colony, particularly at the border (Fig. 2F).

We subsequently analyzed expression of YAP and TAZ that generally play a crucial role for cell-matrix and cell-cell interaction (Totaro et al., 2018). Both transcription coactivators were up-regulated at the edge of the colony (Fig. 2G), which is in line with previous observations (Abagnale et al., 2017). At early time points (at day 3) YAP and TAZ were preferentially localized at nuclear compartment, whereas their localization shifted to the cytoplasm at later time points (day 6; *P* < .0001). However, there was no significant difference in the nuclear/cytoplasmic ratio of YAP and TAZ between edge *versus* center of the colony (Fig. 2H).

To address the relevance of calcium dependent cell-cell interaction for segregation of OCT4 and E-cadherin, we used intermittent calcium depletion by treatment with egtazic acid (EGTA) for 20 minutes at day 5 followed by incubation with rho-associated protein kinase (ROCK) inhibitor-containing medium. Staining of zonula occludens-1 (ZO-1) at day 6 demonstrated that the tight-junctions were not fully reestablished after calcium chelation (Suppl. Fig. S2E). The intermittent calcium depletion in combination with ROCK inhibition - but neither of them individually - abolished the self-organization of OCT4, but not E-cadherin, at the border region of the colonies (Fig. 2I), indicating that cell-cell interaction is relevant for the spatial self-organization.

### Spontaneous formation of 3D aggregates from µCP substrates

After about one week of culture, most of the iPSC colonies detached spontaneously from µCP substrates (Fig 3A; Suppl. Movie S1). While this was initially unintended, we noticed that this detaching worked very consistently without additional enzymatic or mechanical treatment – hence, the method might be applied for generation of early iPSC aggregates. We tested five different iPSC lines and all of them demonstrated reproducible detachment of most colonies within a defined time-window between day 6 and day 8 (Fig. 3B). Within the floating self-detached aggregates, we hardly observed dead cells and they were capable of differentiation into the three germ layers upon induction with FCS-containing medium, in analogy to conventional methods for EB formation (Fig 3C). The size of these self-detaching EBs could be modulated by changing the diameter of the µCP pattern: µCP areas of 400 µm diameter gave rise to aggregates of 248.3 ± 40.6 µm diameter; 800 µm spots produced aggregates of 362.9 ± 51.1 µm in diameter (Fig. 3D). For comparison we also generated EBs with enzymatic harvesting of a cell layer, which resulted in a much larger size-distribution (263 ± 140.4 µm), whereas EB-formation with aggregation of defined cell numbers by centrifugation (Spin-EBs) resulted in smaller variation in aggregate size (Suppl. Fig. S3A,B). We then directly compared the dynamics of differentiation of self-detaching EBs and Spin-EBs over two weeks. Quantitative RT-PCR revealed similar up-regulation of marker genes for the three germ layers (ectoderm: *PAX6* and Nestin; mesoderm: Brachyury and *RUNX1*; endoderm: *SOX17* and *AFP*), and similar down-regulation of *OCT4* (Suppl. Fig. S3C).

**Figure 3:**
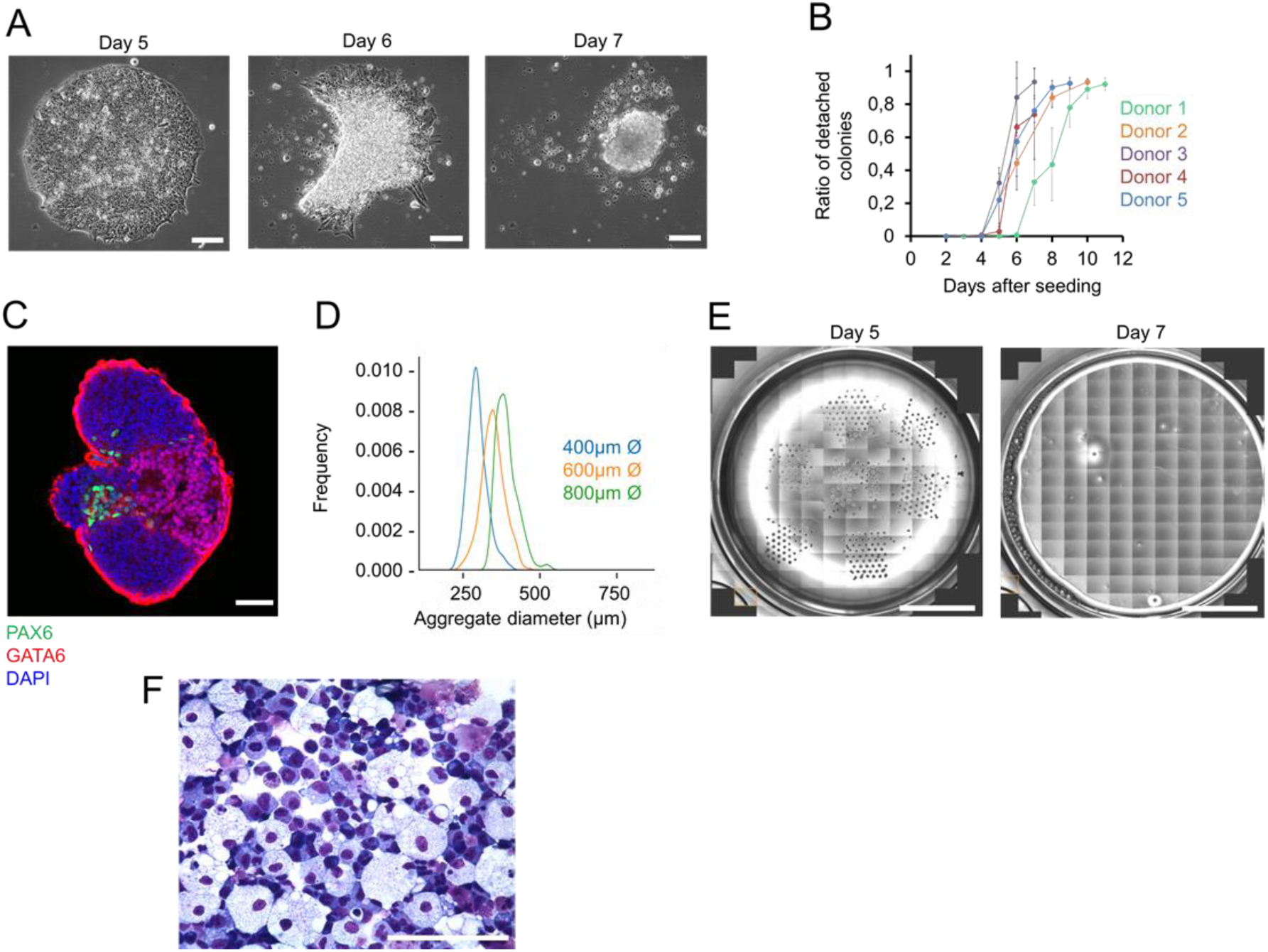
EBs generated by self-detachment from micro-contact printing substrates. **A)** Phase contrast images from a live cell imaging sequence showing the detachment process of a colony from micro-contact printed vitronectin (µCP; Ø 600 µm) on tissue culture plastic. Confluent colonies at day 5 retracted and finally detached at day 7 after seeding (scale bar: 100 µm). **B)**.Kinetics of colony detachment of iPSCs cell lines of five different donors (three replicas for each iPSC lines; ± standard deviation). **C)**.Confocal fluorescence microscopic analysis of a self-detached EB after further differentiation in FCS-containing medium for 7 days. PAX6 expression is indicative for neuroectodermal differentiation, whereas GATA6 is expressed in the definitive endoderm and blocks early epiblast differentiation (scale bar: 100 µm). **D)** Histogram of size-distribution of self-detached aggregates generated with µCP areas of different diameter. **E)** Generation of self-detaching EBs using a liquid handling unit. The phase contrast images depict a whole cell culture well with µCP areas (Ø 600 µm per dot) before (left panel) and after (right panel) the self-detachment process with semiautomated cell culture (scale bar: 10 mm). **F)** Cytospin analysis upon hematopoietic differentiation of semi-automatically generated self-detached EBs reveals typical morphology of various types of myeloid cells (stained with Diff-Quik; scale bar: 50 µm).

An advantage of the self-detaching EBs formation is that the processing steps could be easily automated. In fact, using a liquid handling unit we could reliably harvest uniform EBs (Fig. 3E). These automatically generated self-detaching EBs could also be stimulated for direct differentiation toward hematopoietic lineages in a semiautomatic setting (Fig. 3F, Suppl. Fig. S3D). Taken together, the self-detachment of iPSC colonies from µCP substrates provides a powerful approach to generate size controlled-EBs without the need of enzymatic or mechanical treatment.

### Self-detaching aggregates maintain aspects of the spatial organization

We subsequently analyzed if the spatial self-organization, which we observed in the 2D colonies, progresses upon transition to 3D. After detaching from the µCP substrates, immunofluorescence staining of E-cadherin and OCT4 demonstrated heterogeneity within EBs with marked up-regulation at the outer cell layers. In contrast, Spin-EBs revealed a much more uniform pattern after aggregation (Fig. 4A). Furthermore, in self-detached EBs, the pattern of NODAL expression was very similar to OCT4, having higher expression at the rim than in the center of the aggregates (Fig. 4B).

**Figure 4:**
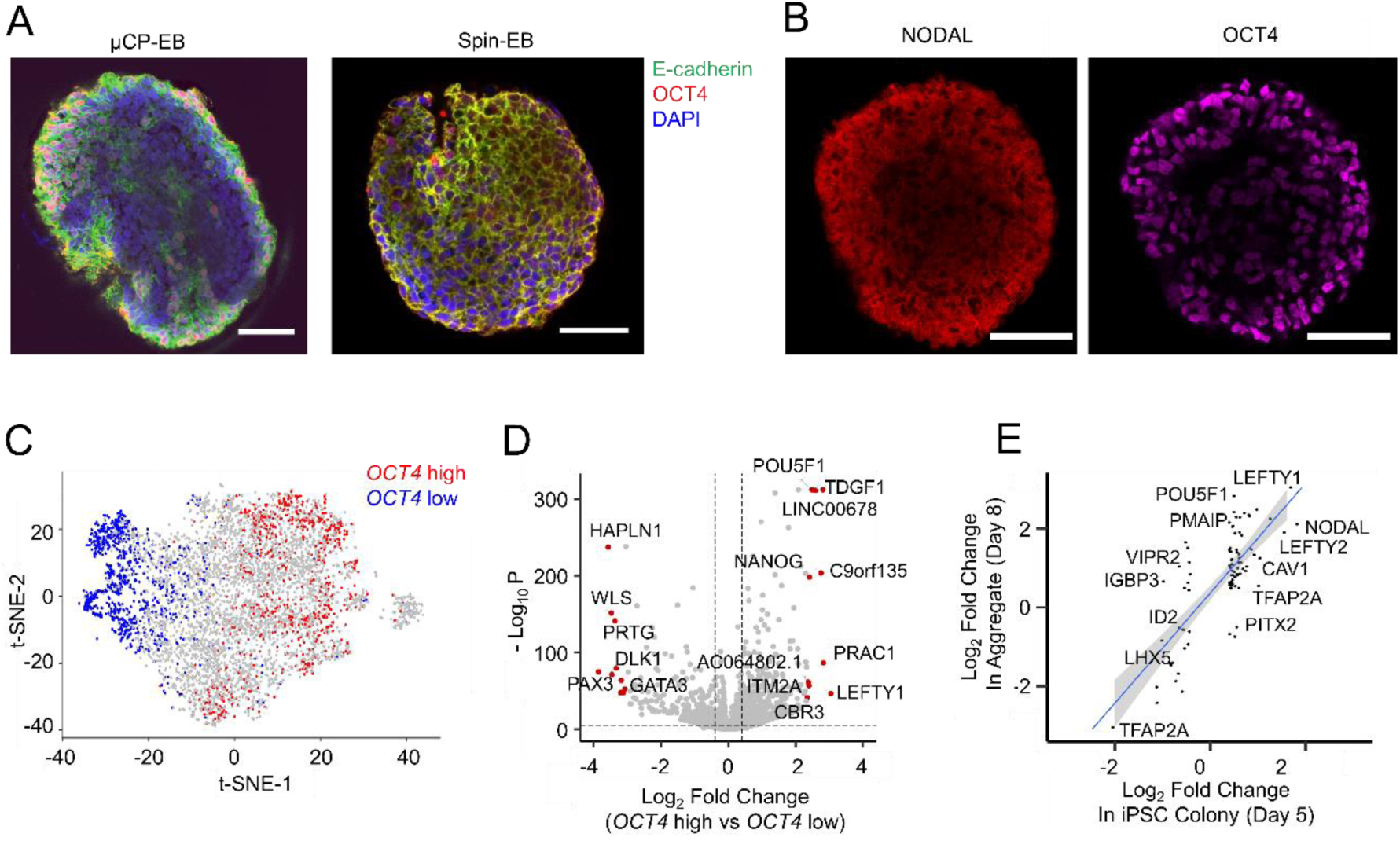
Self-organization within the self-detached 3D aggregates. **A)** Confocal Imaging of aggregates generated by self-detachment from micro-contact printed vitronectin (d8 after seeding of iPSCs on µCP substrates; Ø 600 µm) or pellet formation with centrifugation (Spin-EB; after 24h in same pluripotency-supporting medium; scale bar: 50 µm). **B)** Confocal imaging of self-detached EB reveals that NODAL and OCT4 are higher expressed at the outer layers (scale bar: 50 µm). **C)** Single-cell RNA-sequencing of early self-detached aggregates (d8 after seeding on µCP substrates; Ø 600 µm). The t-SNE plot demonstrates further separation of *OCT4*^high^and *OCT4*^low^subpopulations. **D)** Volcano plot of differential gene expression between *OCT4*^high^ and *OCT4*^low^ subsets. *LEFTY1, NANOG*, and *TDGF1* (NODAL co-receptor) are amongst the genes with highest differential expression. **E)** Comparison of the differential gene expression between *OCT4*^high^ and *OCT4*^low^ within either self-organized iPSC colonies on µCP substrates (day 5 after seeding) or upon self-detachment and early aggregate formation (day 8). Genes that were differentially expressed at both time points are indicated (*R*^*2*^= 0.563, *P* < .001).

To gain insight into the heterogeneity of transcriptional activity within the early self-detaching EBs, we performed scRNA-seq of 3D aggregates at day 8 after seeding of the cells on 600 µm diameter µCP substrates (about 1-2 days after self-detaching of aggregates). We compared gene expression of 1,000 cells with highest *versus* lowest *OCT4* expression in analogy to the differential gene expression analysis in the self-organized 2D iPSC colonies. t-SNE dimensional reduction shows that these two populations were separated more clearly to opposite poles of the t-SNE plot (Fig. 4C). 975 genes revealed significant differential expression between *OCT4*^high^ and *OCT4*^low^ subsets in the aggregates (fold change of ln1.5 and adjusted p-value < 0.05; Suppl. Table S2). Several members of the NODAL signaling pathway (*LEFTY1, TDGF1, NODAL*) were again the highest differentially expressed genes between the two subpopulations (Fig. 4D). To better understand if the differential gene expression in the self-organized 2D colonies progressed from day 5 (in the 2D colonies) to day 8 (in 3D aggregates), we directly compared the fold-changes of *OCT4*^high^/*OCT4*^low^ subsets: overall, the fold changes became even more pronounced in the 3D aggregates at day 8, for both the upregulated and the downregulated genes. Furthermore, genes that were differentially expressed at both time points revealed clear correlation in differential gene expression (Pearson correlation coefficient = 0.755, *P* < .001), indicating progression of the transcriptional heterogeneity (Fig. 4E). The results were validated with self-detached aggregates of another iPSC-line (Suppl. Fig. S4A,B). GSEA indicated that particularly mitochondrial genes, metabolic pathways, and genes of the TGF-ß/SMAD signaling pathway were enriched in the *OCT4*^high^ subset, whereas various developmental pathways were rather activated in the *OCT4*^low^ subset (Supp. Fig. S4C). These results indicate that the cellular specification in the spatially organized 2D iPSC-colonies continue after detaching in the 3D aggregates.

## Discussion

Pluripotency is maintained and stabilized by a network of pluripotency associated genes as well as by external signals (Li and Belmonte, 2017). Previous reports have demonstrated that cells at the edge of PSC colonies have higher expression of pluripotency markers (OCT4, GDF3, Cripto1) as well as higher self-renewal capacity as compared to cells at the colony center (Hough et al., 2009; Hough et al., 2014). Other studies demonstrated that geometric confinement recapitulates the self-organized patterning of human embryonic stem cells (hESCs) (Warmflash et al., 2014). We demonstrate that the spatial self-organization of OCT4 and E-cadherin expressing subsets progresses continuously in geometrically confined iPSCs with culture time. Furthermore, the organization is very similar on PDMS, TCP, and glass, indicating that it is independent of the substrate. Our single cell gene expression analysis demonstrated that the *OCT4*^high^ cells, which are typically localized at the border of the self-organized colony, have similar expression patterns to hESCs subsets that display high self-renewal capacity (Lau et al., 2020). Thus, there is interaction within iPSC colonies that results in spatial self-organization and may ultimately prime cells towards specific cell fates.

What are the mechanisms that govern the developing spatial organization within iPSC colonies? It was striking that amongst the most differentially expressed genes between *OCT4*^high^ and *OCT4*^Low^ cells were *NODAL, LEFTY*, and other components of TGF-ß pathway. This supports the notion that members of the TGF-ß family play a central role in analogy to other self-organization phenomena (Chhabra et al., 2019; Etoc et al., 2016; Warmflash et al., 2014). Other authors used a computational model to simulate the spatiotemporal distribution of signaling components within geometrically confined colonies of differentiating hESCs and indicated that a reaction-diffusion mechanism might govern the complex pattern development during gastrulation (Tewary et al., 2017). Upon establishing of basal-apical polarity of cells, the receptors for TGF-ß ligands, such as receptors for BMP4, NODAL, and TGFß1, are preferentially located at the basolateral domain. This in turn limits the receptor accessibility at the center of the colony resulting in a signaling gradient (Etoc et al., 2016; Nallet-Staub et al., 2015). In fact, we observed that localization of NODAL was most pronounced at the apical surface, especially at border regions of the colony. On the other hand, our scRNA-seq data and immunofluorescence analysis demonstrate that the differential expression in the 2D colonies progresses in the 3D aggregates, accompanied by similar segregation of OCT4^high^ and NODAL^high^ cells at the outer layer. While the mechanism driving the self-organization may differ between 2D colonies and 3D aggregates, the lack of self-organization of Spin-EBs points to the role of pre-organization in 2D colonies in priming the self-segregation of early 3D aggregates.

Direct cell-cell interaction might also be involved in the spatial self-organization (Etoc et al., 2016; Martyn et al., 2019), which is also reflected by the prominent up-regulation of E-cadherin at the border region of colonies. The intermittent disruption of such cell-cell junctions by calcium chelation clearly abolished the self-organization of OCT4, but not of E-cadherin. Recently, a feedback loop of cell-cell contacts - particularly of E-cadherin - and NODAL signaling has been described in mesoendodermal cell-fate specification during zebrafish gastrulation (Barone et al., 2017). A similar positive feedback mechanism may exist between cell-cell contacts and NODAL signaling that primes cells towards specific cell-types already within colonies of iPSCs (Lau et al., 2020). In addition, YAP and TAZ were previously shown to modulate TGF-ß signaling through interaction with SMAD proteins (Ben Mimoun and Mauviel, 2018; Narimatsu et al., 2015; Varelas et al., 2010), but their nuclear/cytoplasmic ratio did not differ at the center *versus* border of the colonies, indicating that contact-inhibition mediated by the hippo pathway does not drive colony segregation.

Furthermore, we describe a new approach to generate EBs from µCP substrates. This method does not require enzymatic treatment or centrifugation steps and can therefore be easily implemented for automated cell-culture regimen. The self-detached aggregates facilitated reliable differentiation into cells of all three germ layers and may therefore provide a viable alternative to conventional methods for EB formation. Notably, at early stages of cell aggregates the self-detached EBs revealed higher self-organization than Spin-EBs, which might be due to their generation from pre-organized iPSC colonies. Albeit the subsequent growth and multilineage differentiation of self-detached EBs and Spin-EBs was very similar, it is conceivable, that there are functional differences that warrant further analysis between different methods for EB formation (Pettinato et al., 2015; Zeevaert et al., 2020).

## Experimental Procedures

### Cell culture of induced pluripotent stem cells

Five human iPSC-lines were generated from mesenchymal stromal cells (iPSC-102, iPSC-104, and iPSC-106) or blood (PT4-WT4, PT18-WT18; Toledo et al., 2020) by reprogramming with episomal plasmids or sendai virus, respectively. The study was approved by the local ethic committee and all samples were taken after written consent (EK206/09). The iPSC lines were cultured on tissue culture plastic coated with vitronectin (0.5 µg/cm^2^) in StemMACS iPS-Brew XF (Miltenyi Biotec). Pluripotency was validated by three lineage differentiation potential and Epi-Pluri-Score analysis, as described in our previous work (Lenz et al., 2015). For intermittent calcium depletion, cell culture medium was aspirated and iPSCs were incubated in 2 mM EGTA (Carl Roth) solution in PBS for 20 mins followed by incubation in culture medium containing Y-27632 ROCK inhibitor (Abcam). Control samples received knockout DMEM (Gibco) for the amount of same time.

### Generation of PDMS pillars

To fabricate an array of PDMS pillars a photomask was manufactured with circular (diameter of 400, 600, 800 µm) and elliptical features (800 x 200 µm, 1200 x 300 µm, 1600 x 400 µm). The patterns were transferred to a silicon substrate using photolithography and were used as a replication mold. PDMS pillars were then constructed using Sylgard 184 (Dow Chemical Company). Pre-polymer and crosslinking agent were mixed at 9:1 ratio, poured over the silicon master, degassed, and cured for 2 hours at 60 °C. Patterned PDMS were then gently peeled-off from the replication mold and were stored in a sealed container until used.

### Micro-contact printing

Micro-contact printing (µCP) was carried out to generate circular adhesion islands with different diameters that facilitate attachment of iPSCs. PDMS pillars were coated with vitronectin (10 µg/mL) for 45 minutes and then used as stamps on pretreated substrates. For quality control, we alternatively used fluorescently labeled gelatin (Molecular probes). Substrates (either tissue culture plastic or glass) were treated with air plasma (50 watts, 1 minute). Stamps were brought into conformal contact with the substrate for at least 1 minute. Patterned substrates were treated using penicillin/streptomycin solution 1:100 overnight, dried, and stored until use.

### Embryoid body formation and differentiation

To form conventional EBs, 80% confluent iPSCs culture were harvested with collagenase IV (Gibco) for 45 minutes and transferred to ultra-low attachment plates (Corning). To form Spin-EBs, iPSCs colonies were treated with Accutase (STEMCELL Technologies) to form single cells. 3,000 cells were dispensed to each well of U-bottom 96 well plates, centrifuged for 6 minutes at 250 RPM, and EBs were harvested 24-48 hours after centrifugation. In contrast, our self-detaching EBs formed spontaneously from micro-contact printed substrates after 6-8 days in culture. Automatic generation of self-detaching EBs was performed on a liquid handling platform (Hamilton Microlab STAR, Hamilton company). Flushing points were determined using whole-well imaging to collect the self-detached colonies.

Multilineage differentiation of EBs was performed with differentiation induction medium (EB-medium) containing Knockout DMEM (Gibco), 20% FCS (Lonza), 2 mM L-Glutamine, 1x Non-Essential Amino acids, 100 nM b-Mercaptoethanol, and 100 U/mL Penicillin/Streptomycin solution. EBs were differentiated in ultra-low attachment plates for 7 days and then transferred to gelatin-coated plates for the reminder of the experiment. Hematopoietic differentiation was carried out as described before (Kovarova and Koller, 2012). Briefly, EBs were incubated in EB-medium in ultra-low attachment plates for 5 days followed by incubation in hematopoietic differentiation medium on gelatin-coated plates with regular medium changes for up to 10 weeks. Hematopoietic progenitors were collected regularly starting from week 3. The cells were analyzed by cytospins (stained using Diff-Quik) and by flow cytometry using a FACS Canto II (BD biosciences) with antibodies for CD31 (Clone: WM59), CD34 (Clone 581), CD43 (Clone:1G10) all BD Bioscience, CD33, CD45 (Clone REA747), CD66b (Clone REA306) from Miltenyi Biotech, and KIT (clone 104D2), CD235a (Clone HIR2) from eBioscience.

### Immunofluorescence

For immunofluorescence analysis of 2D iPSCs colonies, the cells were fixed with paraformaldehyde (PFA) 4% for 10 min, permeabilized with 0.1% Triton X-100 (Carl Roth) for 10 min, and blocked using 2% bovine serum albumin (BSA) for 30 min. Samples were incubated overnight at 4°C with primary antibodies against OCT3/4 (clone: H-134; Santa-Cruz); NANOG (clone: NNG-811), YAP (clone: EP1674Y), and TAZ (all from Abcam); NODAL (Clone: 5C3; Sigma-Aldrich); LEFTY (R&D systems). Secondary antibody staining was done at room temperature for 1-3 hours with donkey anti-goat (Alexa Fluor 488), goat anti-rabbit (Alexa Fluor 594), goat anti-rabbit (Alexa Fluor 647), goat anti-mouse (Alexa Fluor 594), and goat anti-mouse (Alexa Fluor 647); all from Invitrogen. Samples were counterstained with Hoechst 33342 for 10-15 minutes.

The staining of 3D aggregates was done as described before (Dekkers et al., 2019): 3D aggregates were fixed using 4% PFA for 1 hour followed by permeabilization in 0.1% Triton-X 100 and 0.2% BSA (Sigma-Aldrich). Samples were then stained with primary antibody for PAX6 (clone: D-10, Santa-Cruz); GATA-6 (clone: D61E4; Cell Signaling); OCT4, NODAL, and LEFTY for 2 days at 4°C. Secondary antibody staining was carried out for additional 2 days and finally the aggregates were counterstained with Hoechst 33342 for 2 hours. Stained samples were embedded in glycerol-fructose clearing solution and mounted employing a spacer between a coverslip and a glass slide. All samples were imaged using LSM 700 confocal microscopy (Carl Zeiss) using 20x and 63x oil-immersion objective with 2x line averaging or EVOS FL (Thermo Fisher) at 4x, 10x, and 20x. Radial profile of immunofluorescence images of 2D colonies were quantified using radial profile extended ImageJ plugin (Schindelin et al., 2012) and plotted using a custom python script.

### Single cell RNA sequencing

Single cell RNA sequencing (scRNA-seq) was performed with the chromium single-cell gene expression platform (10x Genomics). To this end, we used two iPSC lines that were either geometrically confined at day 5, or within self-detached iPSCs aggregates at day 8 after cell seeding. Colonies and aggregates where treated with Accutase for 15 minutes and sequencing libraries were prepared according to manufacturer’s recommended protocol. Sequencing was done with the Ilumina NextSeq 500 platform and analyzed using Cell Ranger (10x genomics) and the Seurat R package (Stuart et al., 2019). Quality control of the data was carried out using the number of features. After the exclusion of cells with abnormally high/low number of features, at least 6,000 cells per replica were further analyzed. After filtration, average number of features ranged between 2,400 – 3,000 features per cell; average reads were 8,000-10,000 reads per cell. According to the *POU5F1(OCT4)* expression, we classified the top 1,000 cells as *OCT4*^high^ and lowest 1,000 cells as *OCT4*^low^. Differential gene expression between these groups was carried using Seurat wrapper for MAST R package (adjusted p-value of < 0.05 and a threshold of > ln1.5 was considered significant) (McDavid et al., 2015). Gene Set Enrichment Analysis was carried out using Clusterprofiler R package employing fgsea algorithm and gene ontology database (Yu et al., 2012).

### RNA isolation and RT-qPCR

Total RNA was isolated from EBs using Nucleospin RNA kit (Macherey-Nagel) and converted into cDNA using the high capacity cDNA reverse transcription kit (Life Technologies GmbH). Semi-Quantitative PCR (qPCR) was carried out using SYBR green reagent (Applied Biosystems). Primers for the markers of endoderm (*SOX17, AFP*), mesoderm (Brachyury, *RUNX1*), ectoderm (*PAX6*, Nestin), pluripotency (*OCT4*), and housekeeping genes (*GAPDH*) are provided in Suppl. Table S3.

## Supporting information

Mabrouk et al. Suppl Figs S1-S4 and Table S3

Suppl. Movie S1-Time-lapse analysis of self-detaching colony

## Acknowledgments

This work was supported by the German Research Foundation (DFG; WA 1706/8-1, WA 1706/11-1, WA 1706/12-1 and 363055819/GRK2415), the Interdisciplinary Center for Clinical Research (IZKF) within the faculty of Medicine at the RWTH Aachen University (O3-3), and by the Imaging Facility, a core facility of the IZKF within the Faculty of Medicine at RWTH Aachen University. This work was supported in part by the Ministry for Innovation, Science and Research of German Federal State of North Rhine-Westphalia, Duesseldorf, Germany (M.Z.) and by the U. Lehmann donation and CAPES-Alexander von Humboldt postdoctoral fellowship (M.A.S.T).

## Author contributions

M.E.M., R.G., G.A., M.Z., and W.W. were involved in conceptualization of research; V.P, and U.S. were involved in the fabrication of PDMS based patterns; M.E.M, G.A, B.Y, K.Z., Z.M, M.A.S.T carried out the experiments, M.E.M., L.S., R.L, and I.C participated in scRNA-seq analysis, M.E.M and W.W. wrote the manuscript and all authors contributed and approved the final manuscript.

## Conflict of interest statement

W.W. and R.G. are involved in Cygenia GmbH (www.cygenia.com) that may provide service for Epi-Pluri-Score analysis. Apart from this, the authors do not have a conflict of interest.

## Supplemental Material

**Supplemental Figures S1-S4 and Table S3**.**pdf**

Fig. S1: Biomaterials for spatial confinement of iPSC colonies; Fig. S2: Differential gene expression of *OCT4*^high^ *versus OCT4*^low^ cells in self-organized iPSC colonies; Fig. S3: Multilineage differentiation of self-detached embryoid bodies; Fig. S4: Differential gene expression within self-detached aggregates; Table S3: Primer List.

**Suppl. Movie S1: Time-lapse analysis of self-detaching colony**

The phase contrast movie corresponds to the images of Figure 3A. The detachment process of a colony from micro-contact printed vitronectin (µCP; Ø 600 µm) on tissue culture plastic are exemplarily presented. This file is provided as separate avi file.

**Supplemental_Table_S1**.**xlsx**

Table S1: Differentially expressed genes between OCT4^high^ *versus* OCT4^low^ at day 5.

**Supplemental_Table_S2**.**xlsx**

Table S2: Differentially expressed genes between OCT4^high^ *versus* OCT4^low^ at day 8.

