## Supplementary material for "The spatial self-organization within pluripotent stem cell colonies is continued in detaching aggregates": Mabrouk et al. Suppl Figs S1-S4 and Table S3

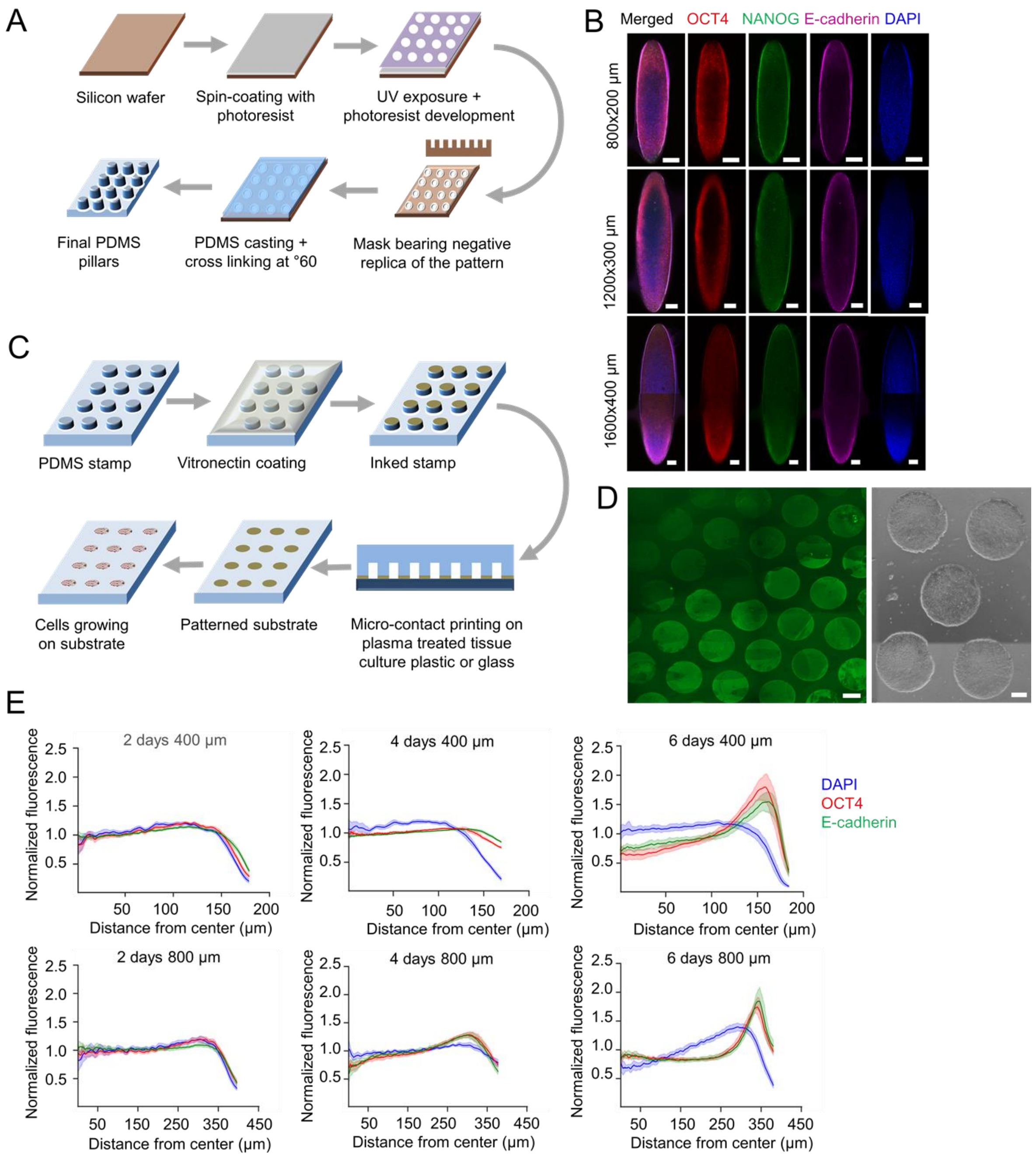


**Suppl. Fig. S1: Biomaterials for spatial confinement of iPSC colonies.**

**A)** Schematic representation of the manufacturing process for PDMS pillars using photolithography.

**B)** Immunofluorescence imaging of spatial organization of iPSCs colonies on top of elliptical PDMS pillars at day 6 (scale bar = 100 µm).

**C)** Schematic representation of the micro-contact printing (µCP) process of vitronectin on tissue culture plastic (TCP) or glass.

**D)** µCP of fluorescent gelatin on TCP substrates (left panel; Ø 400 µm; scale bar = 200 µm) and cells growing to confinement on vitronectin printed on TCP (right panel; Ø 600 µm; scale bar = 300 µm).

**E)** Quantification of the radial profile of immunofluorescence signals within iPSC colonies on µCP substrates (either Ø 400 µm or Ø 800 µm) at day 2, 4, and 6 after seeding (n = 20 per time point).

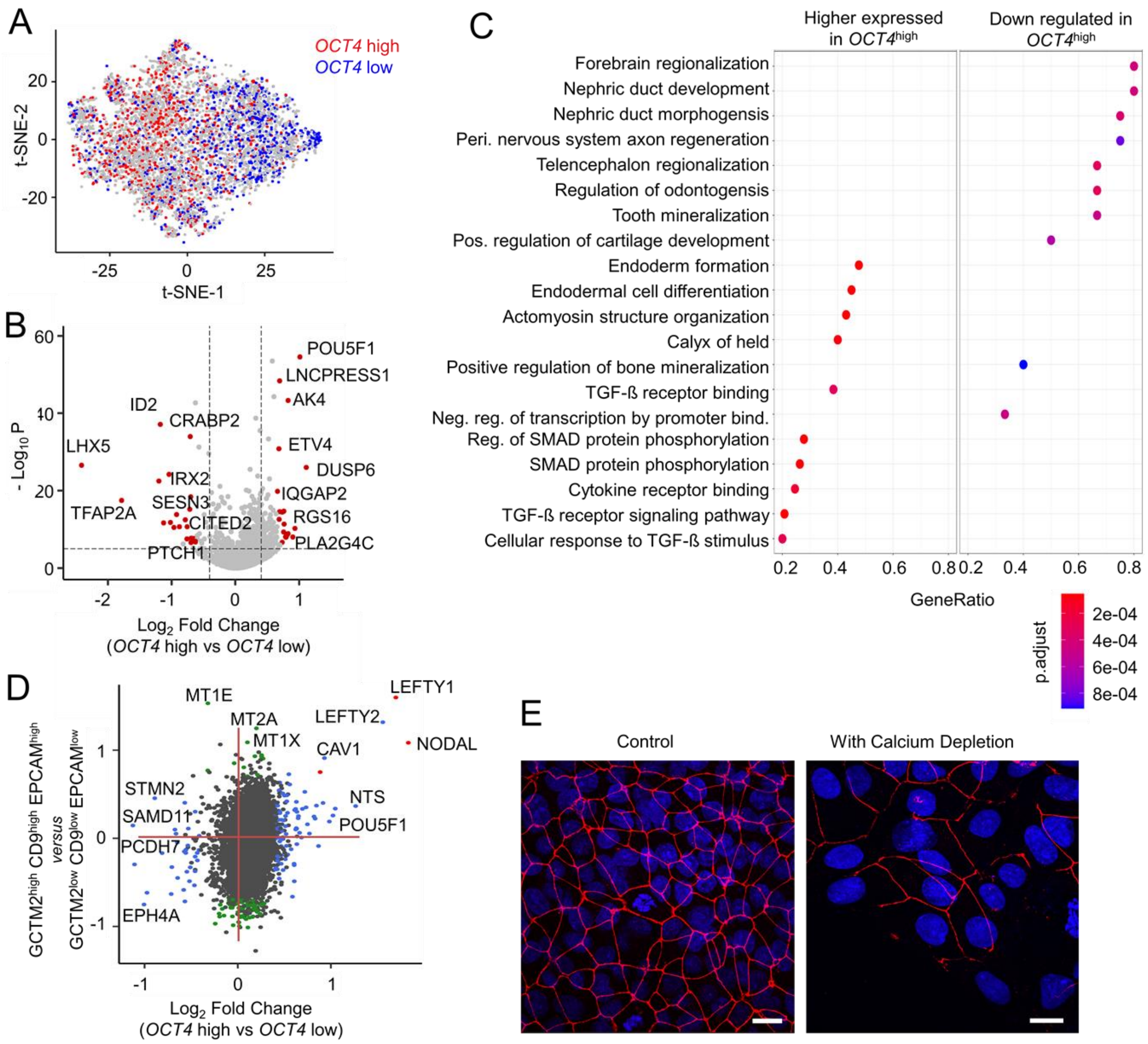


**Suppl. Fig. S2: Differential gene expression of *OCT4*$^{high}$ *versus OCT4*$^{low}$ cells in self-organized iPSC colonies.**

**A)** t-SNE plot of scRNA-seq of spatially confined iPSC colonies on µCP substrates at day 5 after seeding. *OCT4*$^{high}$ and *OCT4*$^{low}$ subsets are indicated. Results correspond to an independent second biological replica (in analogy to Fig. 2A).

**B)** Volcano plot of differential gene expression in *OCT4*$^{high}$ *versus OCT4*$^{low}$ subsets for replica 2 (in analogy to Fig. 2B).

**C)** Gene set enrichment analysis (GSEA) for differential gene expression between *OCT4*$^{high}$ and *OCT4*$^{low}$ cells. Genes in the TGF-β pathway were most significantly overrepresented in the *OCT4*$^{high}$ cells.

**D)** Comparison of differential gene expression between *OCT4*$^{high}$/*OCT4*$^{low}$ subsets ($Log_2$ fold change; replica 1) as compared to differential gene expression between *GCTM2*$^{high}$ *CD9*$^{high}$ *EPCAM*$^{high}$, and *GCTM2*$^{low}$ CD9$^{low}$ *EPCAM*$^{low}$ subpopulations as reported by Lau et al. (2020; Nat Commun doi: 10.1038/s41467-020-16214-8). The *GCTM2*$^{high}$ *CD9*$^{high}$ *EPCAM*$^{high}$ was shown to have higher self-renewal capacity and many genes that were up-regulated in this subset were also higher expressed in our *OCT4*$^{high}$ subset (including *NODAL*, *LEFTY1* and *LEFTY2)*.

**E)** Confocal image of the tight junction marker ZO-1 in spatially confined colonies at day 6. These cells were either cultured without (left) or with intermittent calcium depletion by EGTA for 20 minutes at d5 (right; Scale bar: 20 µm).

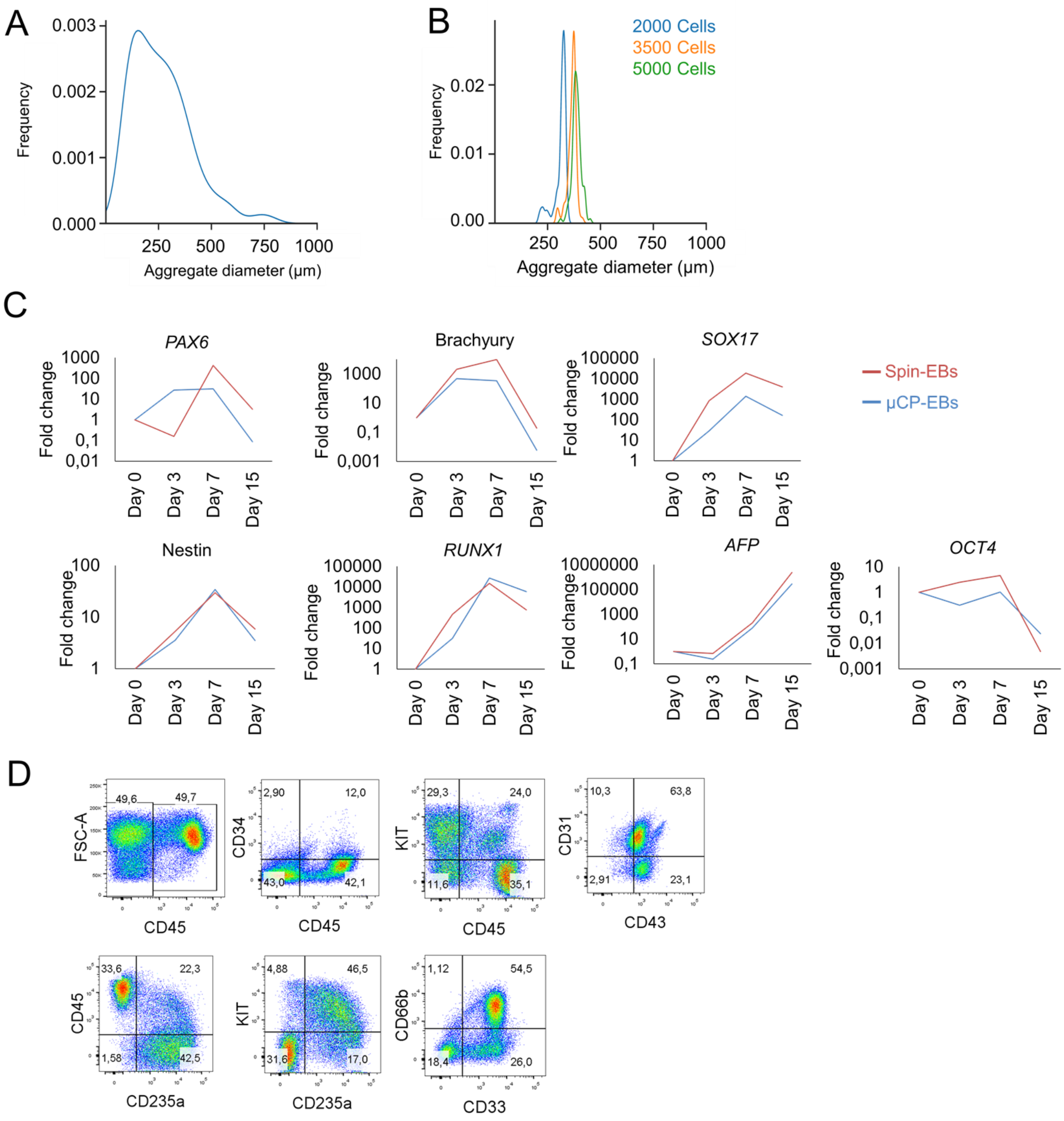


**Suppl. Fig. S3: Multilineage differentiation of self-detached embryoid bodies.**

**A, B)** Histograms of size distribution of early cell aggregates, which were A) generated by collagenase treatment of confluent cell layer of iPSC colonies, or B) by centrifugation of defined cell numbers of iPSCs into pellets (Spin-EB). While aggregate size is much more heterogeneous with the conventional collagenase approach as compared to self-detaching EBs (Figure 3D), it is even more uniform with Spin-EBs.

**C)** Multi-lineage differentiation of Spin-EBs and µCP-EBs after two weeks of differentiation. RT-qPCR analysis of markers for ectoderm (*PAX6,* Nestin), mesoderm (Brachyury*, RUNX1*), endoderm (*Sox17, AFP*), and pluripotency (*OCT4*).

**D)** Flow cytometric analysis demonstrates up-regulation of hematopoietic markers upon hematopoietic differentiation of self-detaching EBs, which were generated semi-automatically using a liquid handling unit.

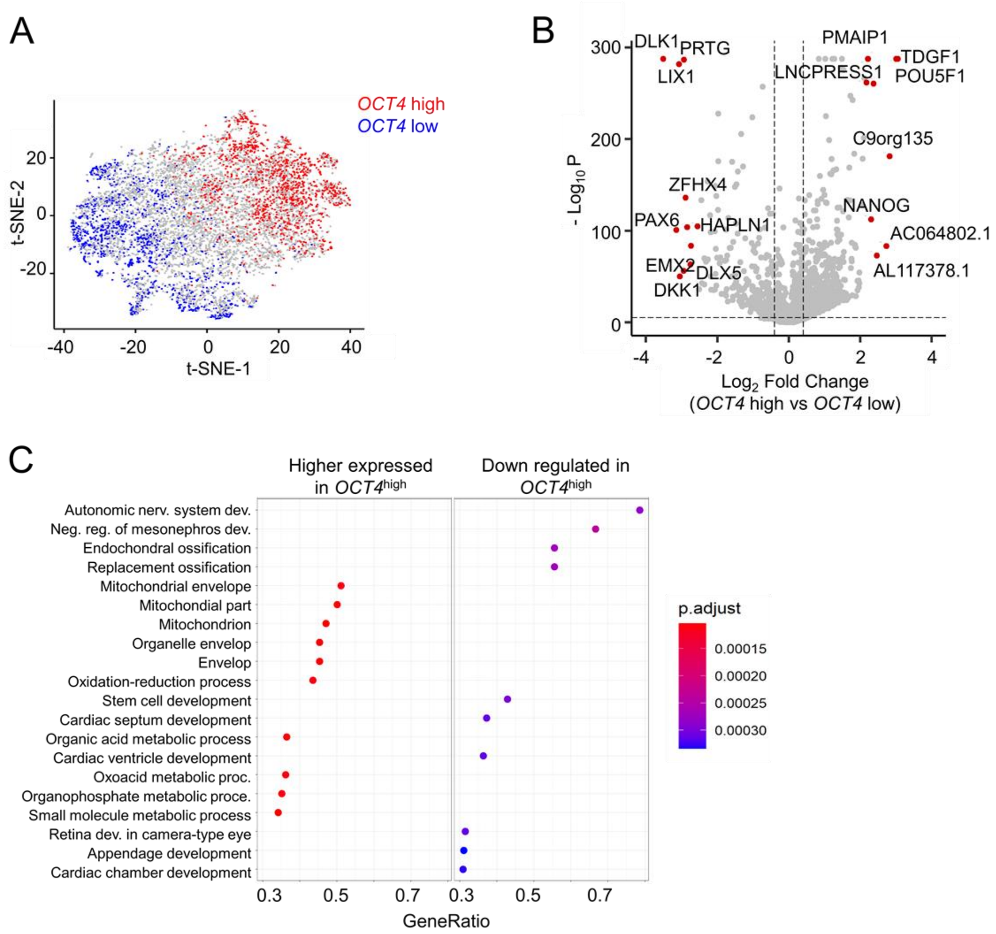


**Suppl. Fig. S4: Differential gene expression within self-detached aggregates.**

**A)** t-SNE plot of scRNA-seq of self-detached aggregate generated with µCP substrates at day 8 after seeding. *OCT4*$^{high}$ and *OCT4*$^{low}$ subsets are indicated. Results correspond to an independent second biological replica (in analogy to Fig. 4C).

**B)** Volcano plot of differential gene expression in *OCT4*$^{high}$ *versus* *OCT4*$^{low}$ subsets for replica 2 (in analogy to Fig. 4D).

**C)** Gene set enrichment analysis (GSEA) for differential gene expression between *OCT4*$^{high}$ and *OCT4*$^{low}$ cells (depicted for replicate 1).

**Suppl. Movie S1: Time-lapse analysis of self-detaching colony.**

The phase contrast movie corresponds to the images of Figure 3A. The detachment process of a colony from micro-contact printed vitronectin (µCP; Ø 600 µm) on tissue culture plastic are exemplarily presented. This file is provided as separate avi file.

**Suppl. Table S1: Differentially expressed genes between OCT4$^{high}$ versus OCT4$^{low}$ at day 5.**

This list of genes, fold-ratios and adjusted p-values is provided as separate EXCEL file.

**Suppl. Table S2: Differentially expressed genes between OCT4$^{high}$ versus OCT4$^{low}$ at day 8.**

This list of genes, fold-ratios and adjusted p-values is provided as separate EXCEL file.

**Suppl. Table S3: Primer List.**

| NAME | DNA SEQUENCE |
|---|---|
| AFP For | GCCAAGCTCAGGGTGTAG |
| AFP Rev | CAATGACAGCCTCAAGTTGT |
| OCT4 For | GGGGGTTCTATTTGGGAAGGTA |
| OCT4 Rev | ACCCACTTCTGCAGCAAGGG |
| PAX6 For | TCGAAGGGCCAAATGGAGAAGAGAAG |
| PAX6 Rev | GGTGGGTTGTGGAATTGGTTGGTAGA |
| Nestin For | CCTCAAGATGTCCCTCAGCC |
| Nestin Rev | CCAGCTTGGGGTCCTGAAAG |
| brachyury For | CAGTGGCAGTCTCAGGTTAAGAAGGA |
| brachyury Rev | CGCTACTGCAGGTGTGAGCAA |
| Runx1 For | CCGAGAACCTCGAAGACATC |
| Runx1 Rev | GTCTGACCCTCATGGCTGT |
| Sox17 For | AGGAAATCCTCAGACTCCTGGGTT |
| Sox17 Rev | CCCAAACTGTTCAAGTGGCAGACA |
| GAPDH For | GAAGGTGAAGGTCGGAGTC |
| GAPDH Rev | GAAGATGGTGATGGGATTTC |
